# A Bidirectional Neural Interface With Direct On-Device Neuromorphic Decoding for Closed-Loop Optogenetics

**DOI:** 10.64898/2026.03.25.714179

**Authors:** G. Bilodeau, A. Miao, G. Gagnon-Turcotte, C. Ethier, B. Gosselin

**Affiliations:** Dept. of Electrical and Computer Eng., Université Laval, Quebec, QC G1V 0A6, Canada; CERVO Research Centre, Quebec, QC G1J 2G3, Canada

**Keywords:** Wireless, Neural Interface, Neural Decoder, Optogenetics, Neuromorphic, Real-Time, FPGA

## Abstract

Bidirectional interfaces combined with neural decoding algorithms are essential for closed-loop (CL) neuromodulation, enabling simultaneous neural monitoring and responsive optogenetic stimulation. However, implementing these capabilities in compact wireless headstages for freely moving animals remains challenging, as most existing platforms rely on tethered setups and external processors to execute computationally intensive decoders. This work presents the design and optimization of a neural decoder integrated into a bidirectional wireless system for CL optogenetic experiments in rodents. The proposed platform combines 32-channel electrophysiological recording with neuromorphic feature extraction, dimensionality reduction, and a nonlinear support vector machine (NL-SVM) decoder implemented on a resource-constrained Spartan-6 FPGA. Temporal dynamics are captured using spike-count features and leaky integrators, while principal component analysis (PCA) reduces the feature space to six components, enabling sub-millisecond inference with minimal memory and power requirements. Model size is further reduced using k-means clustering during training to limit the number of support vectors. Decoder performance was validated using datasets from non-human primate and rat motor cortex recordings. The proposed decoder achieved accuracy comparable to convolutional neural networks (*R*^2^ =0.85 vs. 0.87) and outperformed Wiener filters (*R*^2^ = 0.81) while requiring significantly fewer computational resources. The full system was further demonstrated *in vivo* through wireless closed-loop optogenetic stimulation in rats, achieving a variance accounted for (VAF) of 0.9148. Overall, this work introduces a versatile, fully self-contained, and resource-efficient platform for real-time untethered closed-loop neuroscience experiments.

## I. Introduction

Recent advances in optogenetics have enabled precise, light-based modulation of neuronal activity, reducing artifacts and improving precision compared to traditional electrical stimulation techniques. When combined with electrophysiological recordings, optogenetic stimulation provides a powerful approach for studying neural circuit dynamics, particularly in small rodent models such as freely behaving mice and rats [1], [2], [3].

In parallel, neural interfaces have rapidly advanced with the integration of machine learning (ML) techniques that enable closed-loop (CL) neuromodulation. In these systems, stimulation is guided by features extracted from ongoing neural activity [4], [5], [6], [7], [8]. Such approaches have demonstrated strong potential for studying neural mechanisms and developing treatments for neurological disorders. Implementing closed-loop systems in freely moving rodents with high precision and low latency remains challenging due to hardware constraints, system complexity, and real-time processing limitations. In such paradigms, latency is not only a technical constraint but also a critical biological parameter, as stimulation must remain temporally relevant to ongoing neural activity to effectively influence circuit dynamics and preserve causal relationships. Low-latency stimulation is critical because interventions must remain temporally relevant to ongoing neural activity to effectively influence circuit dynamics and preserve causal relationships. Timing-sensitive plasticity mechanisms, such as spike-timing-dependent plasticity (STDP), typically operate on the order of 10–20 ms [9], highlighting that biologically meaningful modulation requires feedback delivered within this timescale.

The choice of neural biomarkers strongly influences the design and performance of CL neuromodulation systems. Implantable electrodes and microelectrode arrays (MEA) enable simultaneous acquisition of high-frequency signals such as action potentials (APs) and lower-frequency signals like local field potentials (LFPs) [1]. Many CL systems rely on low-frequency biomarkers derived from electroencephalogram (EEG), intracranial EEG, or LFPs to decode motor intent or control stimulation [10], [11]. While successful in applications such as deep brain stimulation for epilepsy and Parkinson’s disease [12], [13], [14], low-frequency biomarkers lack the spatial and temporal resolution required to capture interactions at the level of individual neurons and often introduce additional latency. In contrast, AP-based decoding offers finer spatial precision and has demonstrated improved performance in applications such as post-stroke rehabilitation and movement intent prediction [5], [15]. Nevertheless, AP-based systems face challenges related to channel count, device size, and the need for computationally intensive spike sorting. Real-time processing of high-frequency AP signals therefore requires fast and power-efficient hardware, particularly as channel counts increase.

Despite these requirements, many existing systems rely on tethered connections to external computers for signal decoding and stimulation control [4], [16], [17], employ simple or inflexible decoding algorithms [18], [19], or use custom ASICs that lack versatility and are rarely integrated into wireless *in vivo* experimental setups [20], [21], [22]. Consequently, there remains a strong need for compact, wireless, and adaptable CL neural interfaces capable of high-resolution recording, efficient processing, and precise stimulation.

A practical compromise used in many brain–machine interface (BMI) systems is the use of unsorted action potentials or threshold crossings, commonly referred to as multi-unit activity (MUA), as biomarkers [23], [24]. This approach leverages population activity across tens to hundreds of channels and can effectively decode behavioral intent when combined with ML algorithms [25], [26], [27]. However, many such systems rely on deep neural network architectures such as long shortterm memory (LSTM) networks, which require large training datasets and significant computational resources [28], [29], [30], [31]. As a result, these decoders often depend on wired connections to external computers or graphics processing units (GPU), limiting their applicability in freely moving animal experiments.

Although several neural decoder ASICs have been proposed [21], [32], they frequently rely on low-frequency biomarkers or require large FPGA platforms connected to PCs, making them impractical for small, mobile subjects. This highlights the need for a lightweight, wireless, and resource-efficient neural decoding platform.

To address these challenges, we propose a wireless hybrid-neuromorphic neural headstage capable of real-time closed-loop decoding and stimulation in freely behaving rodents. The system emphasizes computational efficiency through neu-romorphic feature extraction, dimensionality reduction, and model optimization, enabling neural decoding within the strict power, memory, and processing constraints of a compact FPGA-based platform.

The decoder is implemented on a resource-limited Spartan-6 FPGA using an optimized nonlinear support vector machine (NL-SVM) model designed to reduce memory and computational requirements while maintaining high decoding performance. The FPGA implementation enables sub-millisecond decoding and supports simultaneous wireless datalogging of neural data for live experimental validation and analysis. The decoder output can be used to trigger closed-loop optogenetic stimulation. The proposed system was validated using neural datasets from motor tasks recorded in both primates and rodents, demonstrating reliable neural decoding performance. Closed-loop capabilities were further demonstrated in an *in vivo* experiment with anesthetized rats, where cortical multi-unit activity recorded from a microelectrode array was used to wirelessly trigger optogenetic stimulation of dopaminergic neurons in the ventral tegmental area (VTA).

Overall, this work presents a compact wireless bidirectional neural interface that integrates efficient neural decoding and real-time optogenetic stimulation within a single untethered platform. By enabling low-latency neural decoding, reconfigurable decoding model and stimulation, and simultaneous wireless data logging, the proposed system provides a flexible tool for advanced behavioral neuroscience experiments in small freely moving animal models.

## II. System Overview

The wireless headstages material consists of 4 main blocs [33]. The neural recording interface, the optical stimulation, the wireless transceiver and the digital signal processing (DSP) and control of the device.

The RHD2132 from *Intan Technologies* is used for neuro-recording. It can record differential signal from 32 output with one common reference, with a maximum amplitude of -5mV to 5mV and a sampling rate of up to 30 ksps. The total recording bandwidth is set by the chips bandpass filter, which can be from 0.1 Hz to 20 kHz. The chip measures only 8×8 mm.

The optical stimulation circuit is based on the AS1109 8 channels, 8-bit light-emitting diode (LED) driver. Each channel of the chip can supply from 0.5 to 100 mA. On the system, each LED is driven by two channels in parallel configured identically allowing each LED to get a maximum of 200 mA current. For example, at a current of 150 mA, using high power LEDs such as: LB G6SP from OSRAM Opto Semiconductors, the system can deliver 250 mW/mm^2^ at the output of an optical fiber. The chip is in a 16 pins QFN package and measures only 4×4 mm.

The wireless transceiver is the nRF24l01p from *Nordic Semiconductor*. It is a 2.4GHz transceiver using a custom protocol by *Nordic Semiconductor* with a maximum on air data rate of 2 Mbps, a 11.3 mA current in transfer mode and a 13.5 mA in receive mode at 2 Mbps. The chip is in a 20 pins QFN package and measures only 4×4 mm.

In this design, an FPGA is utilized to perform all of the DSP and classification. This includes AP detection and wavelet AP compression using dedicated custom core modules developed previously by our group [34]. A novel hybrid-neuromorphic CL core is also integrated in this module. Within this CL core, a new fully integrated, real-time neuromorphic feature extraction and reduction core is integrated which extracts the AP firing rate calculation into bins, calculates a leaky integrator for each channel and infers PCA score for the first 6 principal components. The second part of the CL core is the NL-SVM, which can perform classification and regression tasks in order to control optical stimulation based on the neural activity.This feature extraction and neural decoding chain is shown in Figure 1.

**Fig. 1.**
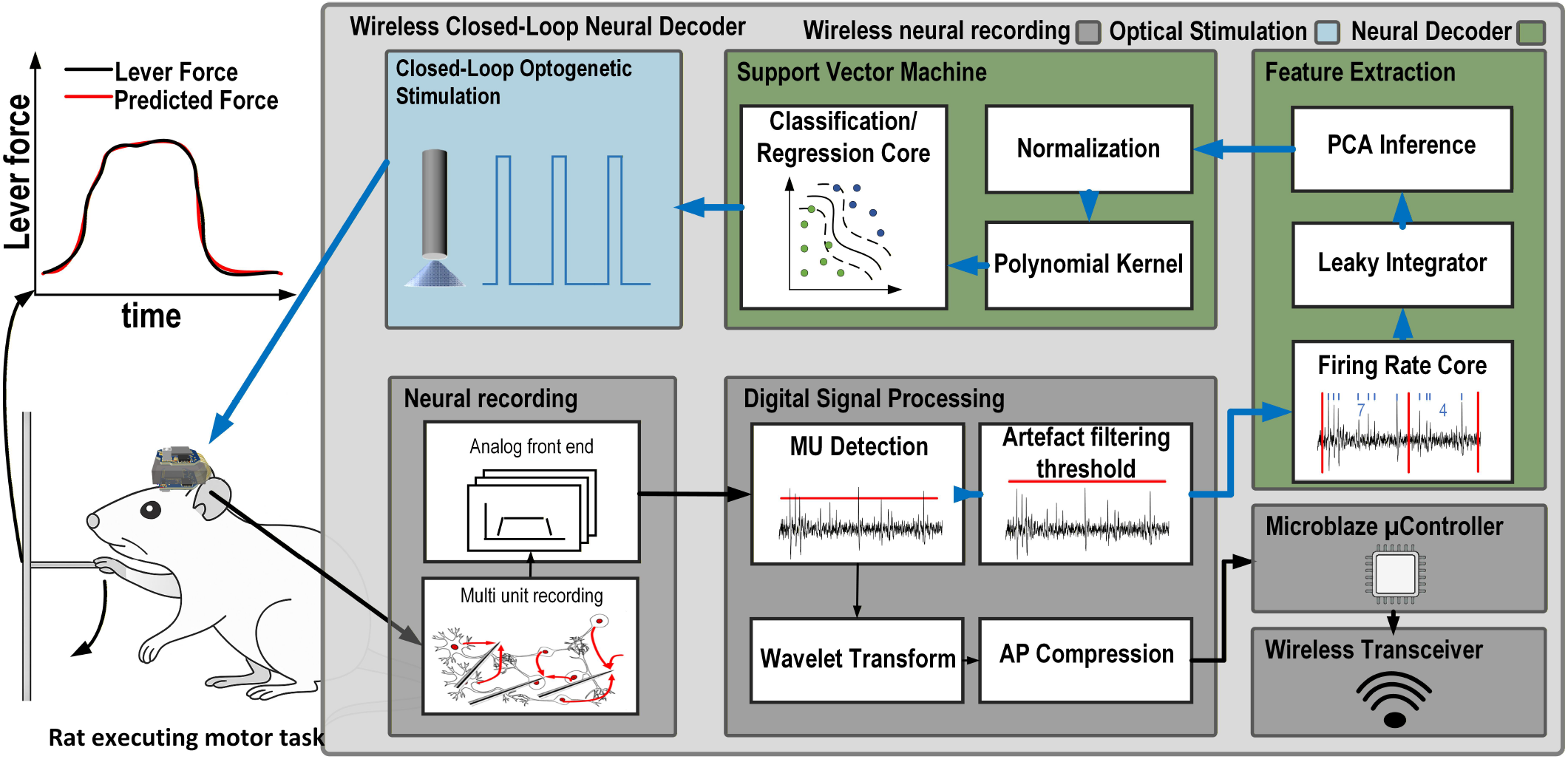
Block diagram and overall concept of the wireless headstage and neural decoder. Left: A rat performs a lever-pull task while carrying the wireless headstage. The force applied to the lever can be predicted in real time by the onboard neural decoder using an SVM-based regressor. Right: Block diagram of the decoding pipeline, showing the neural interface, digital signal processing stages, binned spike counts, leaky-integrator filtering, PCA-based feature extraction, and the nonlinear SVM decoder. The decoder output can be used to drive closed-loop optical stimulation.

The FPGA controls all the custom digital cores and all the other integrated circuits (ICs) using a *Microblaze* microcontroller unit (MCU) soft core, e.g. the neural interface, the wireless transceiver and the optical stimulator. The block diagram of the system is depicted in Figure 1.

Figure 2(a) shows the unfolded headstage leveraging a thin rigid-flex printed circuit board with all components identified. Since the system is designed for experiments on small size rodents, it should minimize weight while not compromising performance and autonomy (30-60 minutes). This system weight 4.68 g with a 100 mAh battery (Model 051417, MYD Technology)) and including the plastic packaging and on/off switch (∼ 1.7 g w/o battery, 3.8 g w/battery only). With this packaging and battery, the system is suited for experiments with bigger laboratory mice or rats.

**Fig. 2.**
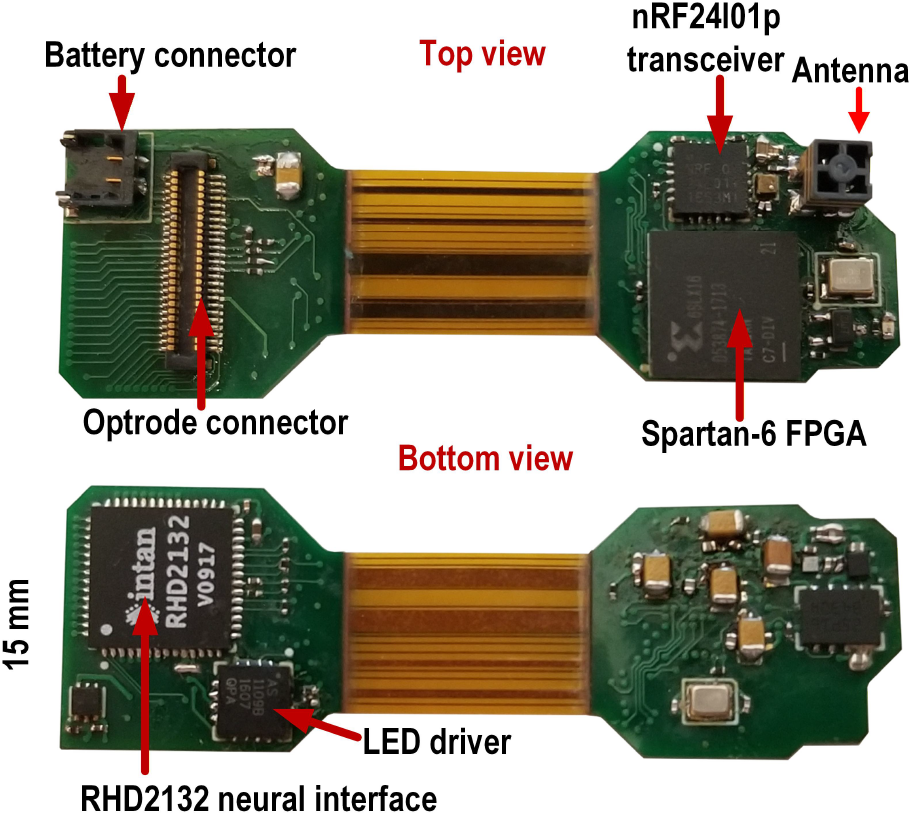
Two-sided view of the rigid-flex PCB when not folded [33]

While the printed circuit board (PCB) and the AP detection and compression cores were previously presented in [34], the novel real-time CL neural decoding core featuring the integrated feature extraction and NL-SVM cores, along with the experimental analysis and *in vivo* CL experiments presented are the main focus of this paper.

## III. Decoder Model Overview

In order to integrate a neural decoder into a resource-constrained system, both memory and computational demands must be significantly optimized. In this work, two primary strategies are employed to make such integration feasible: *feature optimization* through neuromorphic feature extraction and *model optimization*.

### A. Feature Optimization

High-performance neural decoders often treat neural signals as time-series data [35], where the current signal state depends on previous states, much like in speech processing. Traditional classifiers such as CNNs [35] and SVMs typically incorporate multiple temporal bins of history, sometimes up to 300 ms. In LSTM-based decoders, this temporal context is naturally maintained. However, including temporal history usually increases the number of features significantly, thereby raising the computational and memory load.

To address this issue, a neuromorphic architecture is adopted to capture the temporal dynamics of the neural signal while reducing feature size. Event-based MUA detection and accumulation, combined with a brain-inspired *leaky integrator*, condense the contribution of multiple time bins into a single feature representation [29], [36]. The leaky integrator assigns higher weight to more recent events while gradually discounting older activity. This approach allows temporal information to be preserved without explicitly maintaining a large temporal history.

Using the leaky integrator greatly reduces the decoder’s input size and computational demand compared to traditional decoders. For example, a decoder with 32 channels and a history of 20 time bins per channel would produce 640 features in a conventional implementation, whereas the leaky integrator produces only 32 features. Previous methods have also employed a boxcar filter combined with an SVM classifier to incorporate multiple time bins and improve classification accuracy [29]. However, this approach significantly increases detection latency, up to 900 ms in some cases [29], because neural activity over the time window is averaged with equal weight. In contrast, the leaky integrator assigns greater weight to more recent data, reducing feature size, improving accuracy, and decreasing decoding latency.

Further dimensionality reduction is achieved by applying PCA after leaky integration. Retaining only the top six principal components results in an 81% reduction in feature dimensionality relative to the 32-channel feature vector. This reduction allows the system to focus on the most informative aspects of the data by projecting channel activity onto a compact set of orthogonal components.

The choice of retaining six principal components is based on a trade-off between predictive performance and FPGA resource constraints. Performance was evaluated using data recorded from a rhesus macaque performing a center-out isometric wrist force task, previously reported in [37]. The effect of reducing the number of retained principal components on prediction performance (measured using *R*^2^) is shown in Figure 3. Both predicted x- and y-axis forces exhibit a clear elbow around six components, beyond which additional components yield only marginal improvements in accuracy (*<* 2%).

**Fig. 3.**
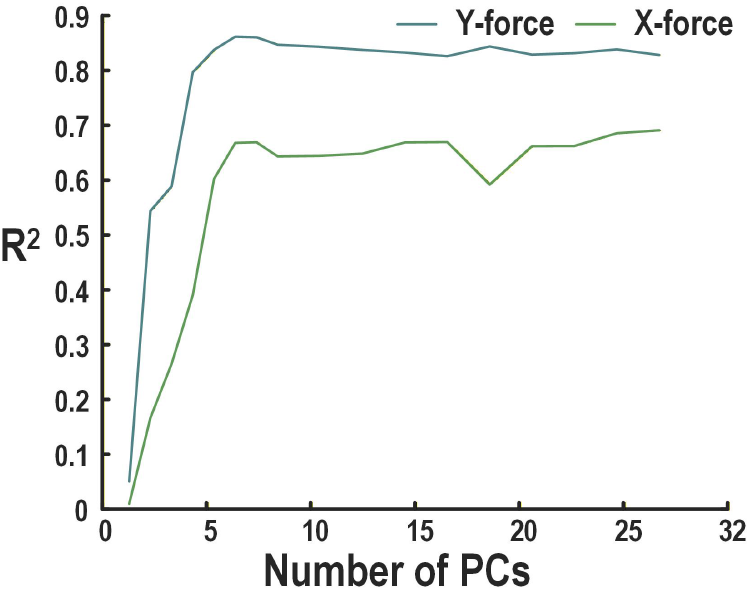
*R*^2^ for different number of principal components kept when predicting the x and y-forces using the dataset from the center-out isometric wrist force task from [37] using an SVM decoder

Since the number of features directly impacts FPGA memory usage and computational cost, the smallest value within the plateau region (i.e., the elbow point) was selected. The overall model size scales linearly with both the number of input features and the number of support vectors in the SVM. Therefore, reducing feature dimensionality allows more resources to be allocated to the SVM model, improving flexibility under fixed hardware constraints.

Although the optimal PCA dimensionality is inherently dataset-dependent, the selected value reflects the intrinsic dimensionality observed in representative training data and provides a stable operating point for deployment. While a larger number of principal components could be retained if required under different conditions, this would reduce the memory available for the SVM model and increase inference cost on the FPGA.

### B. Model Selection and Optimization

SVMs are highly adaptable due to the *kernel trick*, which enables them to model nonlinear relationships and makes them suitable for a wide range of decoding tasks. However, the often used Gaussian kernel, while powerful, introduces a significant computational burden. To balance versatility and efficiency, this work employs a *polynomial kernel*. This kernel is advantageous in low-power systems because it can be implemented using a single multiply-accumulate (MAC) operation and allows multiple kernel configurations to be supported using the same hardware architecture.

Although the polynomial kernel reduces computational complexity compared to the Gaussian kernel, nonlinear SVMs still require considerable memory to store support vectors. To reduce this memory requirement, a combination of the *reduced set method* and *K-means clustering* is used. This approach groups similar feature vectors in the training dataset and generates a synthetic representation for each cluster. A new, smaller model is then trained on this synthesized dataset to recover accuracy lost during the reduction process. This training procedure is illustrated in Figure 4(a).

**Fig. 4.**
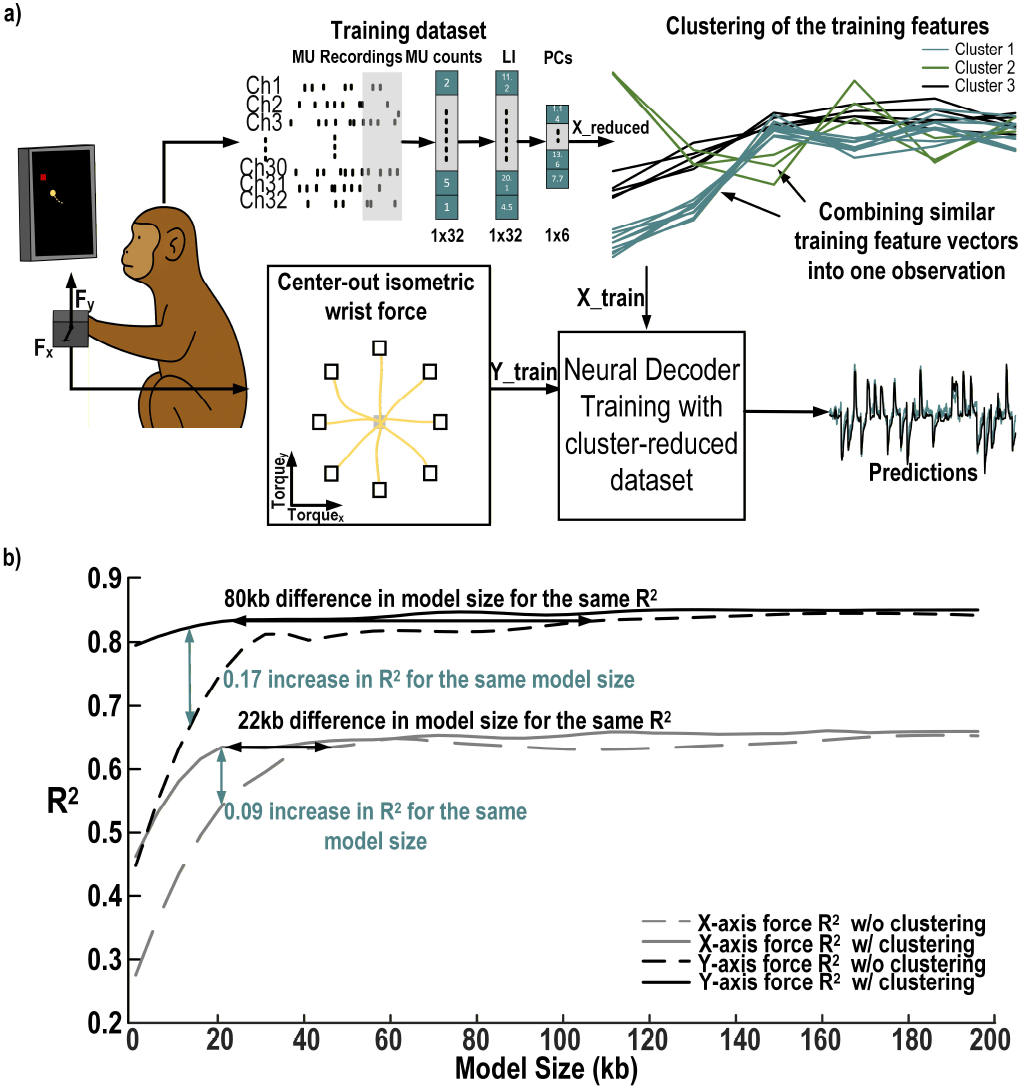
Training process of the proposed neural decoder and its model reduction performance with, a) the cluster based data reduction, from left to right, the monkey performs a behavioral task, while MUA is recorded in the motor cortex to form a training dataset for the SVM algorithm. This data set is clustered using k-means to reduce the size of the training dataset and thus reduce the SVM model size. The SVM is trained on this new, reduced dataset to predict the X and Y wrist forces during a center out isometric wrist force task b) its performance predicting the X and Y-axis forces applied by a monkey performing a center out task from 32 channels of MUA. Comparison of the *R*^2^ values for the X and Y predictions as a function of training size based on training dataset size w/and w/o clustering.

Together, these optimizations produce a compact model that fits within the memory constraints of embedded systems while maintaining performance close to that of the original model, with only a small decrease in decoding accuracy.

## IV. Feature Extraction Core

The feature extraction core is separated in four main steps:

1. The AP detection core, extracting the APs or Multi-Unit activity from the bandpass filtered activity,
2. The event-based Firing Rate Core, which calculates the firing rate for each channel into customizable time bins, computing features only when an event occurs,
3. The brain-inspired Leaky integrator core, which transforms the 32 channels of binned firing rate into brain inspired time dependent features,
4. The PCA inference core, which reduces the 32 channels of leaky integrator results into the 6 components with the maximum explained variance.

Each of these cores are detailed in the following sections.

### A. AP Detection Core

In order to extract the APs from the signal, an adaptable threshold is computed based on a calculation of the standard deviation of the signal. This automatic threshold was previously presented [33] to improve overall AP detection rate and reduce false positives from movement artefacts. The threshold can also be fixed to customizable values on the fly for versatility of the wireless system.

### B. Firing Rate core

The first feature extracted from the detected APs is the firing rate. This feature represents the level of unsorted neural activity observed on each channel within a fixed time window. In this system, the time window is user-defined and can range from 5 to 50 ms, allowing the architecture to support multiple applications. Larger window sizes are typically used for motor decoding tasks [15], while smaller windows provide lower latency and capture neural dynamics on a smaller time scale. This is useful for applications such as AP-based closed-loop epilepsy treatment [5]. It is also beneficial for applications that require stimulation with small delays to leverage spike-timing-dependent plasticity (STDP), where the precise timing of spikes controls synaptic strengthening or weakening [38], [39].

To compute the firing rate for each of the 32 MUA channels, a custom VHDL binned MUA threshold-crossing counter core was developed. This core accumulates the number of threshold crossings detected on each channel during a user-defined time window. The core receives detected events from the AP detection core as input and applies an upper threshold to filter out potential large artifacts, as commonly done in MUA-based neural decoding systems [15] to improve feature quality. When an event successfully passes the upper-threshold filtering step, the firing rate core increments the count associated with the corresponding channel. Once the time window duration is reached, the firing rates for all channels are passed to the leaky integrator core as input features, and all counts are reset to zero.

Timing is managed using the global neural interface timer, which has a resolution of 50 *µ*s. This timer generates a signal to advance the time window at the user-defined interval. Each counter is stored using 4 bits. The design also leverages event-driven computation, allowing the core to remain active only during MUA threshold crossings and window transitions, thereby improving overall power efficiency.

### C. Leaky Integrator core

The leaky integrator core constitutes the second feature extraction step. This step allows the current feature to integrate information from past firing-rate states, with more recent values weighted more strongly than older ones. As a result, only 32 leaky integrator values (one per channel) are passed to subsequent processing stages, representing a significant reduction in feature dimensionality.

The VHDL core that processes the 32-channel data consists of two MAC operators, a 64 × 16 distributed RAM block storing decay coefficients, and a state machine controlling memory access. The discrete leaky integrator equation implemented is:

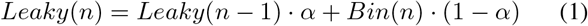

where *α* is the leaky integrator decay, Bin(n) is the current bin count for a specific channel, and Leaky(n-1) is the previous state of the leaky integrator.

The leaky integrator decay is given by:

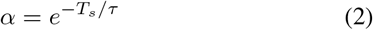

where *T*_*s*_ is the sampling period (bin size) and *τ* is the desired time constant. Each channel’s decay can be customized and is stored using 16 bits in the distributed RAM. Values of 1 −*α* are also precomputed and stored to reduce on-board computation. The leaky integrator computation for each channel is performed in two steps. First, Leaky(n-1) · *α* is computed using the MAC operator’s multiply unit. Second, Bin(n) · (1 - *α*) is passed to the MAC operator and added to the first product using the MAC operator’s accumulation unit. This calculation is repeated sequentially for all 32 channels. Time multiplexing the computation reduces the total hardware required. The latency for one channel is only 7 clock cycles; therefore, the total latency added by time multiplexing is small (225 cycles, or 11.25 µs at 20 MHz) compared to implementing each channel in parallel. Figure 5 illustrates this core.

**Fig. 5.**
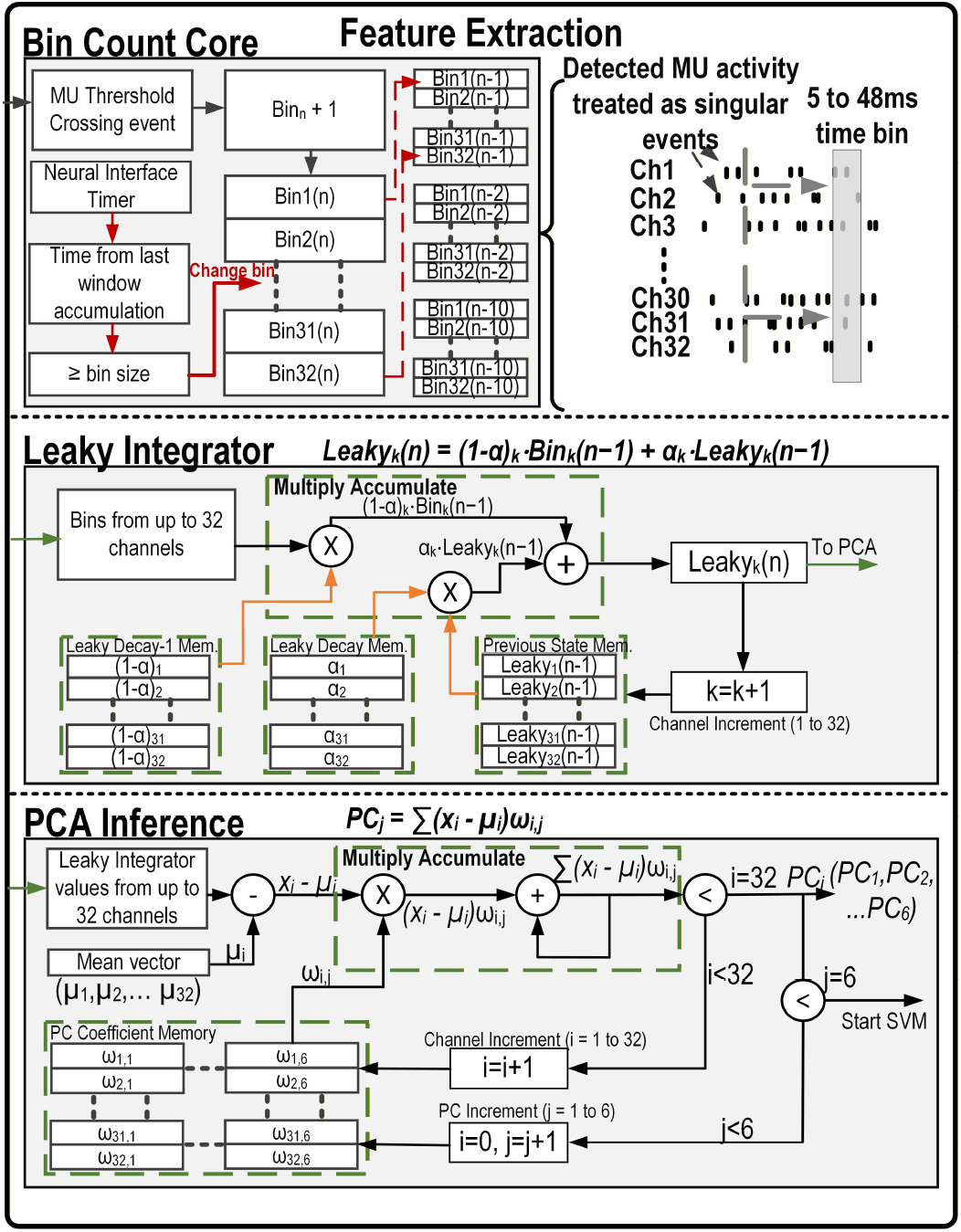
Detailed VHDL implementation of the feature extraction cores with the bin count core (top) that detects MUA threshold crossings, applies an artifact removal threshold and keeps track of the MUA events for each of the 32 channels during a customization time window (bin). A trigger is sent along with the MUA threshold crossing count for all 32 channels to the Leaky Integrator core (middle) when the time widow exceeds the user defined time. The leaky integrator core then calculates the new leaky integrator value for each channel using the user defined coefficients loaded and previous LI state into RAM and a multiply accumulate calculation block. When the 32 LI values are ready, a trigger is sent to the PCA inference core (bottom), where the LI values and the user defined PCA coefficients are used to calculate the coefficients of the 6 principal components at this time point. These 6 values are the features used in the SVM decoder.

### D. Principal Component Analysis Inference Core

The final feature extraction step aims to further reduce the size of the feature vector. During decoder training, a PCA is computed on the recorded 32-channel leaky integrator values to identify the principal components that capture the most variance. This step reduces the feature dimensionality, allowing a smaller set of components to represent the original 32 channels. The trained feature means and PCA coefficients for each principal component are then used to compute the corresponding values for new incoming data in real time.

When applied to new data for dimensionality reduction from a feature vector of size *m* to *k* principal components, the PCA algorithm follows these steps:

1. Standardize the dataset.
2. Compute the *m ×m* covariance matrix.
3. Compute the eigenvectors and eigenvalues.
4. Select the *k* eigenvectors corresponding to the largest eigenvalues.
5. Transform the data using these *k* eigenvectors.

In this work, only the six eigenvectors with the largest eigenvalues are retained and loaded for on-device inference. Computing the PCA transform on incoming data is equivalent to calculating the dot product between each eigenvector (principal component coefficient vector) and the feature vector at a given time:

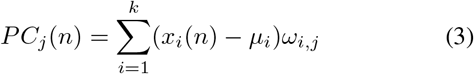

where *PC*_*j*_(*n*) is the *j*^*th*^ principal component at time *n*, k is the number of features (32 here), *x*_*i*_(*n*) is the input feature for channel *i, µ*_*i*_ is the mean of feature *i* computed during training, and *ω*_*i,j*_ is the *i*^*th*^ element of the *j*^*th*^ eigenvector.

On the FPGA, this computation is implemented using a single MAC operator that is time-multiplexed to sequentially compute the six principal components. First, the feature means are subtracted from the incoming feature vector using the MAC operator and pre-loaded −*µ*_*i*_ coefficients. Each adjusted feature value is then multiplied by its corresponding eigenvector coefficient and accumulated to compute the *j*^*th*^ principal component. Once all six principal components are computed, they are passed to the SVM for classification or further dimensionality reduction. The entire computation requires 1047 clock cycles, corresponding to 52.35 µs with the system clock.

## V. SVM C ore

In order to make predictions based on the features extracted from the MUA while minimizing feedback latency, an onboard support vector machine inference core is implemented. This SVM performs real-time predictions with sub-millisecond delays using the previously described features. The following sections detail the FPGA architecture of the proposed SVM core.

### A. Polynomial Kernel Core

The kernel trick allows a linear SVM to learn non-linear patterns by implicitly mapping input data to a higher-dimensional space without explicitly performing the transformation. In this implementation, a polynomial kernel is used to approximate this mapping, enabling the SVM to classify non-linear data using a linear decision function. The kernel core is implemented on the FPGA using a time-shared MAC unit to minimize hardware resource usage. A state machine sequentially feeds the support vectors, stored in block RAM, to the MAC unit to compute the kernel function:

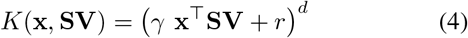

where **x** is the input feature vector, **SV** are the support vectors of the model, *r* is the bias term, *γ* is the scale factor of the polynomial, and *d* is the degree of the polynomial.

Different polynomial degrees are supported by looping the MAC operation and feeding its output back as input for each iteration, effectively computing the power *d*. This design makes the kernel core easily configurable across models with varying polynomial degrees, enhancing the flexibility of the SVM decoder. The full implementation is illustrated in Figure 6.

**Fig. 6.**
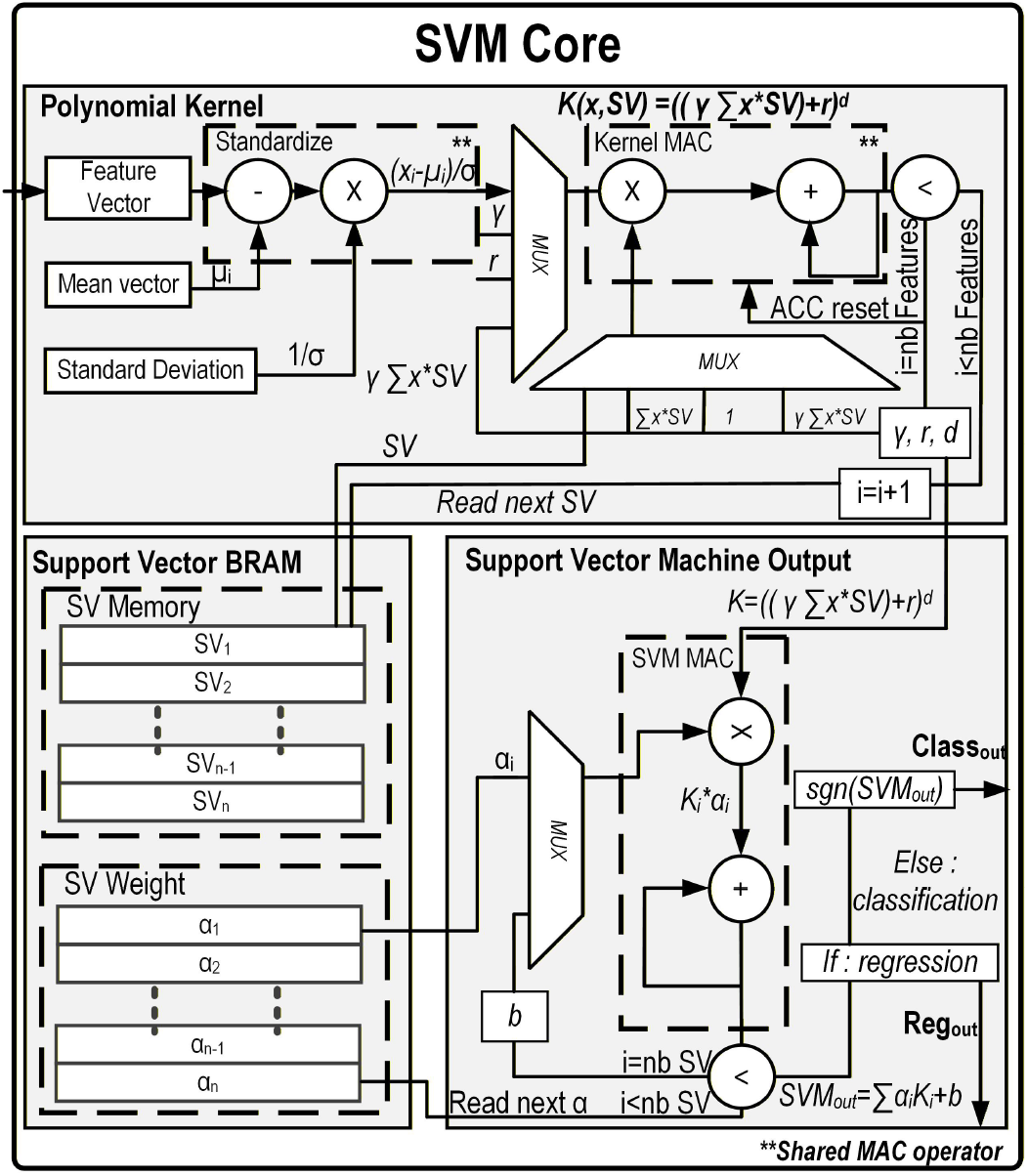
Detailed VHDL implementation of the SVM core with a polynomial kernel. The design includes three main components: the polynomial-kernel block, the SVM output block, and the reprogrammable block RAM storing the support vectors (SVs) and their weights. Top: The polynomial-kernel module receives each stored SV and incoming feature vector, applies standardization based on the trained parameters, and computes the *d*^*th*^-degree polynomial kernel using a time-multiplexed MAC operator. Bottom right: Once each kernel value is generated, it is forwarded to the SVM output core, which accumulates the weighted kernel contributions incrementally. After all SVs have been processed, the final output is produced, either as a continuous value for regression or as a sign-based decision for classification.

### B. Real-Time FPGA SVM Core

The real-time FPGA SVM core is also shown in Figure 6. The kernel outputs are combined with their corresponding weights (*α* values), which are then used to compute the linear SVM decision function. The decoder can be configured either as a classifier, outputting binary labels corresponding to decoded movements, or as a regressor producing continuous output trajectories.

The maximum memory usage for the SVM model is 10.5 kB, and a complete inference requires up to 3805 clock cycles (190.2 *µ*s with the 20 MHz system clock).

### C. Wirelessly Programmable Model

For freely moving experiments in small rodents, the neural decoder model must be versatile and reprogrammable on-the-fly. During a startup sequence, the desired model can be transmitted wirelessly to program the FPGA RAM through the communication chip. This approach is highly flexible and allows the user to configure:

- Firing-rate bin size, from 5 ms to 50 ms,
- Leaky integrator decay parameters, independently for each of the 32 channels,
- PCA coefficients, which can be retrained to fit specific datasets,
- SVM model and support vectors, with options for linear, second-order polynomial, and third-order polynomial kernels,
- Closed-loop stimulation trigger parameters, including threshold, hold time, and rest period,
- Stimulation parameters for a pulse-width modulation (PWM) pattern, such as delivered current, frequency, and duty cycle.

## VI. Measured performances and results

### A. Feature optimization Results

To evaluate the proposed feature optimization and assess the potential reduction in overall model size, the performance of several decoders using different feature extraction methods was compared, as shown in Table I. The dataset used for these experiments was recorded from a rhesus macaque performing a center-out isometric wrist force task, previously reported in [37]. A subset of 32 electrodes (channels 33 to 64) was selected to emulate the constraints of the wireless decoder proposed in this work.

**Table I.** Comparison of accuracy for different features and decoders, highlighting the feature size of each case.

| Feature | Feature Size | Model | Y force $R^2$ | X force $R^2$ |
| --- | --- | --- | --- | --- |
| Binned MUA | 32 | Linear Regression | 0.32 | 0.17 |
|  |  | NL-SVM | 0.50 | 0.34 |
|  |  | CNN | 0.54 | 0.30 |
| 10 Bins of History | 320 | Linear Regression | 0.73 | 0.50 |
|  |  | NL-SVM | 0.85 | 0.71 |
|  |  | CNN | 0.87 | 0.70 |
| Leaky Integrator | 32 | Linear Regression | 0.77 | 0.57 |
|  |  | NL-SVM | **0.84** | **0.71** |
|  |  | FFNN | 0.85 | 0.73 |
| Leaky Integrator + PCA | 6 | Linear Regression | 0.77 | 0.44 |
|  |  | NL-SVM | **0.85** | **0.66** |
|  |  | FFNN | 0.86 | 0.68 |

The decoder aims to predict the X- and Y-axis forces generated during the task. The recorded force signals were used to segment the dataset and provide ground truth for training and testing the decoding algorithms. Raw neural signals were thresholded to detect spike events, which were then accumulated into 50 ms bins to compute the firing rate for each channel. Depending on the feature configuration being evaluated, a leaky integrator was subsequently computed for each channel following (1) with a decay time constant corresponding to a 300 ms period. PCA was then applied to the resulting feature vectors using the training data to compute the principal component coefficients. The dataset was divided into 70% training data and 30% testing data, preserving temporal divide in data in order with timeseries predictions. The CNN used in testing consisted of three 2D convolutional layers with ReLU activations and batch normalization, followed by a fully connected layer with 128 units and dropout, and a linear output layer that predicts force from sequences of 32 neurons over 10 time bins. A feedforward neural network (FFNN) is also used to compared performance when features are extracted prior to training using LI and PCA. The SVM uses a second order polynomial kernel. The performance of the different feature sets and classifiers is summarized in Table I. This comparison highlights both the feature dimensionality and decoding accuracy for each configuration, demonstrating the effectiveness of the leaky integrator combined with PCA in maintaining high decoding accuracy while significantly reducing the feature vector size.

*The resources for the proposed method include the wireless reprogramming logic and full logic to compute feature extraction for 32 channels.

### B. Model Optimization Results

To evaluate the potential model size reduction achieved with the SVM clustering optimization, the dataset described in Section VI-A was used. Models were trained to predict the X- and Y-axis forces, with the dataset split into 70% for training and 30% for testing. SVM models were trained using different training dataset sizes. In the baseline condition (without clustering), training samples were randomly selected from the training dataset. In the clustering condition, the training vectors were grouped into a specified number of clusters to approximate the full dataset using fewer representative samples. This approach reduces the number of support vectors in the trained SVM, resulting in a smaller model.

The results, compared with SVMs trained on randomly selected subsets of the original training data, are shown in Figure 4(b). These results demonstrate a potential reduction in model size between 22 kb and 80 kb for an equivalent

*R*^2^ when minimizing model size without sacrificing decoding accuracy. Conversely, when the model size is constrained to 15–20 kb, clustering improves *R*^2^ by 0.09 to 0.17 compared with random sampling.

These results show that clustering is an effective strategy for reducing the SVM model size while maintaining decoding performance, allowing the model to fit within the memory constraints of the headstage FPGA while minimizing hardware footprint and power consumption.

### C. Cross Species Prediction Results

To further demonstrate the proposed architecture and its ability to generalize across species and neural decoding tasks, the model was evaluated on two datasets: the monkey dataset described in Section VI-A, recorded during a center-out isometric wrist force task, and a dataset recorded from a rat performing a lever-pull task.

1. *Monkey dataset prediction results:* Using the dataset, feature extraction pipeline, and training method described in Sections VI-A and VI-B, the neural decoder achieved prediction accuracies (*R*^2^) of 0.85 and 0.66 for the Y-axis and X-axis forces, respectively.
2. *Rat dataset classification results:* The dataset used for this validation consisted of recordings from a 16-channel electrode array implanted in the motor cortex of a rat performing a lever-pull task. In this task, the rat was required to pull a lever and maintain force production for up to two seconds. The applied force was measured using the MotoTrak behavioral module (Vulintus, Inc.) [43]. A trial was initiated once a force of 10 g was exerted on the lever. The rat then had two seconds to reach a target force to receive a food pellet reward.

The target force varied between 15 g and 200 g and was adaptively adjusted based on the animal’s recent performance. Specifically, the target force increased slightly when the success rate exceeded 70% over the previous 10 trials and decreased when it fell below 50%. On average, the target force was approximately 100 g, with a success rate ranging between 50% and 60%.

To validate the decoder, a binary classification task was defined to predict when the rat started pulling the lever above the 10 g initiation threshold. Samples exceeding this threshold were labeled as *pull*, whereas samples below the threshold were labeled as *no pull*. Recordings from different days were evaluated separately, with the model retrained to accommodate changes in the electrode signals, reflecting typical usage scenarios. The dataset was split into 70% training and 30% testing sets, preserving the temporal order to better reflect real-world conditions. The classifiers evaluated include an LDA, a second-order polynomial kernel SVM, a FFNN, and a CNN with architectures similar to those described in Section VI-B. The classification performance, measured in terms of accuracy, macro F1 score, macro precision, and macro recall, across multiple recording days is presented in Figure 9.

As the *pull* class is significantly underrepresented compared to the *no pull* class, misclassification costs were adjusted to achieve a balance between overall accuracy and minority class performance in terms of precision and recall. While the CNN showed slightly improved performance on certain days, its results remained comparable to those of the other established classifiers (LDA,FFNN) and to those of the proposed feature extraction and SVM pipeline as shown in Figure 9.

These results demonstrate that the proposed feature ex-traction pipeline and model architecture generalize well to classification tasks and to neural data recorded in rat subjects.

### D. Neural Decoder Footprint

The VHDL implementation of the optimized neural decoder results in a compact, low-footprint, and low-power solution that can be integrated into a small on-headstage FPGA. The overall resource usage of the neural decoder is summarized in Table II, along with a comparison to other neural decoder implementations. Compared with existing devices, this implementation combines low memory usage with low hardware resource utilization to minimize overall footprint and power consumption. The system was implemented and validated *in vivo* within a wireless, reprogrammable platform.

**TABLE II.**
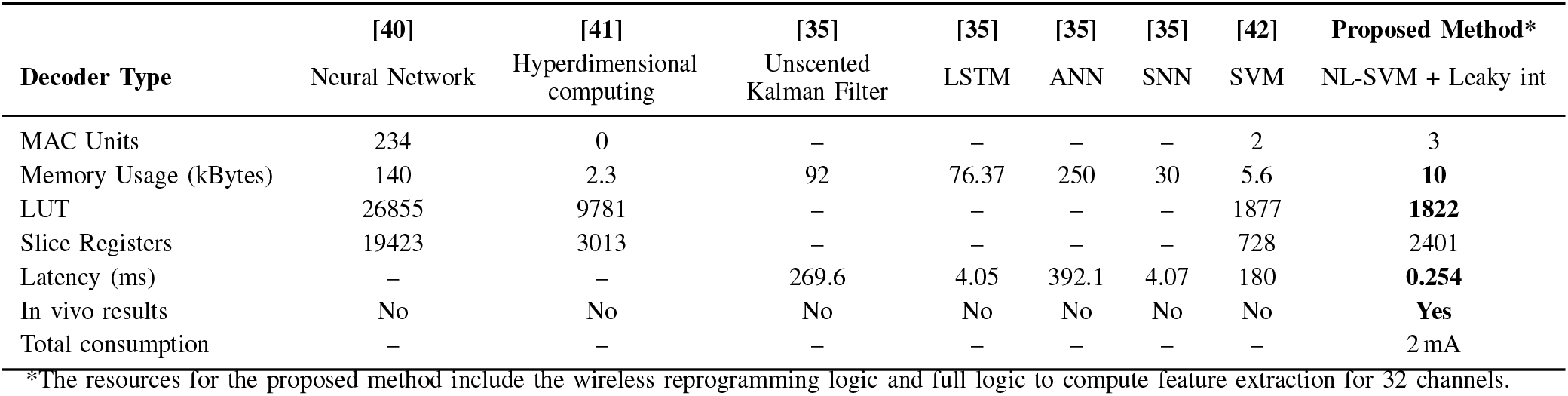
Resource usage and performance comparison of different neural decoders.

Figure 7 shows the percentage of power consumed by each block of the system, including the portion allocated to the neural decoder module and the contributions of each of its main computational cores.

**Fig. 7.**
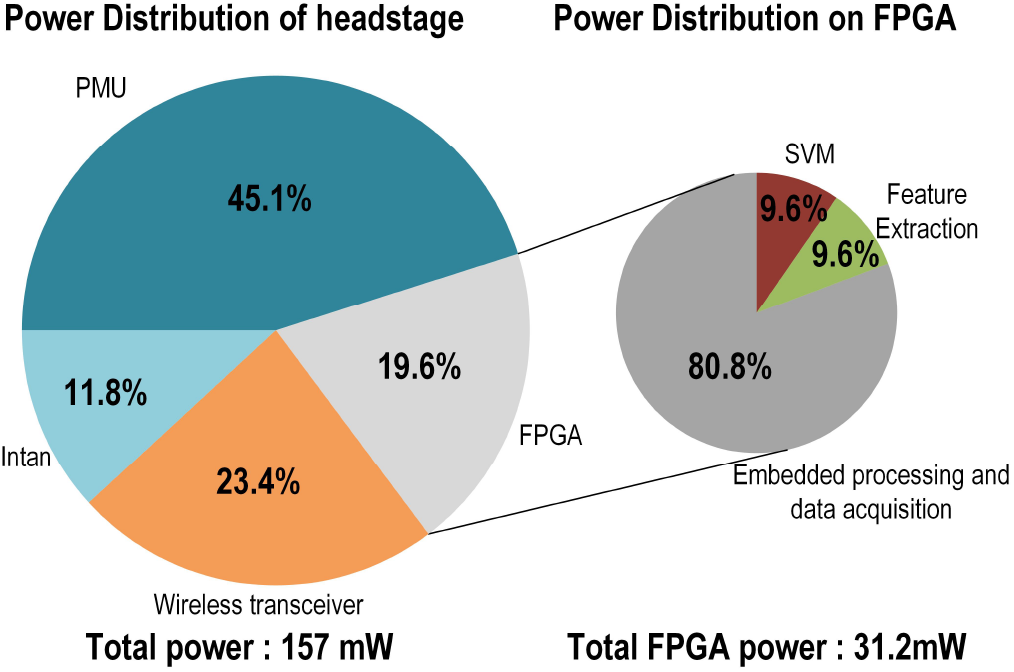
Pie chart of the headstage power consumption per module and a close up on the usage of the neural decoder building blocks implemented on the FPGA.

### E. Real-Time Wireless Neural Decoding Results

To validate the neural decoder on the wireless headstage and demonstrate the reprogrammability of the leaky integrator, PCA, and SVM models, the system was tested using synthetic input signals. A simple model was trained on the data recorded by the headstage to decode global neural population activity. The thresholds, leaky integrator, PCA parameters and the SVM model were sent to the headstage through BLE in a startup sequence. During testing, the decoder output was transmitted in real time along with the detected and compressed APs. The embedded decoder performances were then assessed by comparison with a computer-based calculation of population activity and a computer based decoder operating on global activity extracted from the headstage recordings.

To emulate usage in an *in vivo* setting, a Tektronix AFG3101 arbitrary waveform generator was used to output a custom waveform derived from prerecorded neural signals. This output was connected to all channels of the headstage through a custom-made PCB that divides the input voltage by a factor of 1000 so that it remains within the voltage rating of the Intan chip. The headstage was connected to its adapter through an Omnetics-to-Molex adapter. The experimental setup is shown in Figure 8(c).

**Fig. 8.**
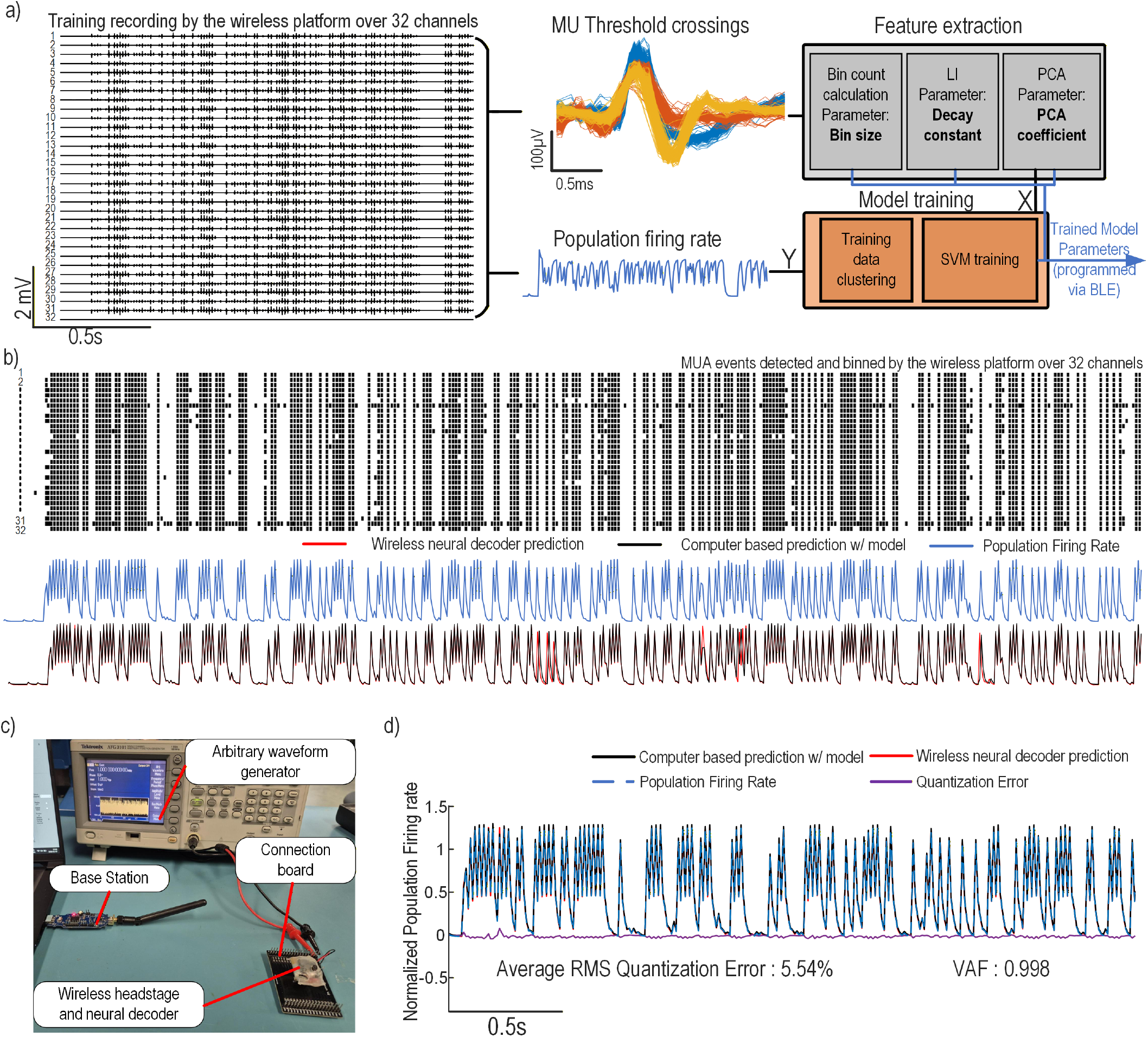
Wireless neural decoder headstage: experimental results and validation with synthetic data played from an arbitrary waveform generator. a) Model training workflow. On the left, a 32-channel synthetic recording with overlaid MUA threshold crossings shows signal capture and detection. Timestamps are used to compute spike-count bins, optimize the LI decay constant, and extract PCA coefficients (gray). Features and population firing rate are clustered to reduce model size, and an SV regressor is trained to predict firing rate. Feature parameters and the trained SV model are transferred via BLE at startup. Recorded activity along with the neural decoder prediction are then transferred wirelessly to the base station in subsequent experiments. b) Wireless decoding results. Top: raster plot of detected MUA across channels. Bottom: population firing rate, on-device prediction, and computer-based prediction from logged MUA (pre-quantized model). c) Experimental setup: headstage, Molex-to-Omnetics adapter, custom signal-injection board, and signal generator replaying prerecorded activity. d) Decoder output and quantization error: VAF = 0.998 and mean RMS error = 5.54%, demonstrating high on-device fidelity.

A recording of the synthetic signal was first performed on the 32 channels to generate a training dataset. To validate the system’s ability to perform fast, real-time neural decoding predictions in a reprogrammable manner, a model was trained based on this recording to approximate the population firing rate from the detected neural activity across the 32 channels. The trained SVM model uses 10 ms bins to accumulate neural activity, a leaky integrator time constant of 10 ms for all channels, and the six principal components extracted from the leaky integrator outputs as inputs to the SVM decoder. The input neural data and target output signal were then processed using the training data clustering algorithm to reduce the model size before training the SVM model on the resulting subset. This process is illustrated in Figure 8(a).

To validate the reprogrammability and functionality of the wireless decoder headstage, the trained model was transmitted via Bluetooth during a startup sequence. The wireless head-stage then recorded APs on 32 channels while simultaneously performing SVM-based neural decoding predictions. The resulting raster plot of the detected APs and the neural decoding output from the wireless decoder is shown in Figure 8(b), along with the predictions obtained from a computer-based implementation of the model and the actual population firing rate.

A closer comparison of the predicted signal from the wireless headstage, the computer-based model, and the actual population firing rate is shown in Figure 8(d). The quantization error between the wireless neural decoder prediction and the computer-based model is also presented. The resulting average RMS quantization error is 5.54%, while the variance accounted for (VAF) of the headstage prediction reaches 0.998. These results demonstrate that the wireless neural decoder can record, detect, and bin MUA activity while performing configurable leaky integrator, PCA, and SVM inference directly on the headstage.

## VII. *In vivo* EXPERIMENT

In this paper, we performed *in vivo* validation of our wireless neural decoding and closed-loop optogenetic stimulation platform. Experiments were conducted in anesthetized rats to simultaneously record motor cortex (M1) activity and deliver closed-loop stimulation to the VTA, enabling realtime evaluation of system performance. Our primary objective was to establish proof-of-concept by demonstrating the ability to acquire multi-channel neural signals, generate on-device predictions of population firing rate, and trigger stimulation with precise timing.

VTA stimulation provides an indirect means of modulating cortical activity. Unlike direct cortical stimulation, it does not deterministically drive the recorded neurons. Specifically, optogenetic activation of VTA dopaminergic neurons induces dopamine release, which acts as a reinforcement signal that modulates excitability and biases plasticity in downstream circuits, including M1. This mechanism increases the likelihood of specific activity patterns rather than directly evoking them, and has been shown to drive the progressive reshaping and reinforcement of M1 activity [44]. Closed-loop optogenetic approaches have further demonstrated that real-time, state-dependent stimulation can causally modulate cortical dynamics [45], [46]. In this context, low-latency operation is critical, as the effectiveness of reinforcement depends on precise temporal alignment between detected neural activity and stimulation. This enables selective and adaptive reinforcement of targeted activity.

Based on these principles, we expected VTA stimulation to induce measurable modulation of M1 activity through this neuromodulatory mechanism. Together, these experiments validate both the functionality of the system and its potential for causal, feedback-driven modulation of cortical circuits.

All animal manipulations were performed with the approval of Universite Laval’s Animal Protection Committee (CPAUL) Protocol # 2023-1256.

### A. Injection surgery and viral vectors

Two long-evens rats (500g male) received injections of AAV2/9-CaMKIIa-ChroME2f-mRuby (Canadian Optogenetic and Vectorology Foundry, 1x6E12 gc/ml) into the left ventral tegmental area (VTA; AP: -5.4, ML: 0.5, DV: 7.2 and 7.5 mm from bregma, 300 nl each) during a stereotaxic surgery under anesthesia. At the same time, a fiber-optic cannula with an integrated LED (525 nm, 400-µm core, NA 0.66, Doric Lenses Inc.[47]) was implanted above the left VTA (AP: -5.4, ML: 0.5, DV: 7.0 mm from bregma).

### B. Intracortical Electrode Arrays Implants and Recording Sessions

Four to six weeks after viral injection, two rats were implanted with intracortical electrode arrays during a second stereotaxic surgery. Two types of implants were used. The first was a polymer array provided by the Polymer Implantable Electrode Foundry at the University of Southern California (BRAIN Initiative Award U24NS113647) [48]. The second was a 32-channel, 4-shank silicon probe provided by the NSF MINT Hub at the University of Michigan (Award 1707316) [49]. Electrodes were targeted to the left motor cortex (AP: 0–4 mm, ML: 1–4 mm, DV: 1.5 mm from Bregma). All experiments were performed intraoperatively under ketamine– xylazine anesthesia.

The custom wireless system was used to perform simultaneous optogenetic stimulation of the VTA and cortical recordings. During these experiments, neural signals recorded by the headstage were transmitted wirelessly to a base station. The wireless headstage also generated drove the LED in the implanted optical cannula. The experimental setup is illustrated in Figure 10(a).

**Fig. 9.**
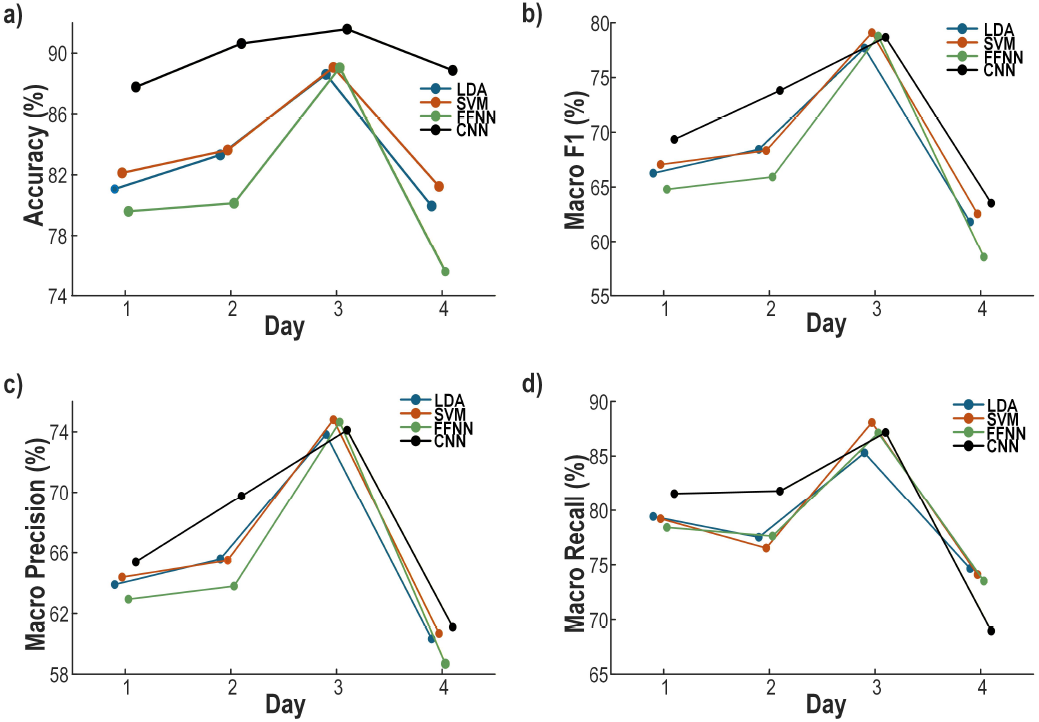
Classification performance comparison of multiple classifiers on the rat lever-pulling dataset across four recording days. a) Accuracy of the classifiers across the four days. b) Macro F1 score over the same period. c) Macro precision over the same period. d) Macro recall over the same period.

**Fig. 10.**
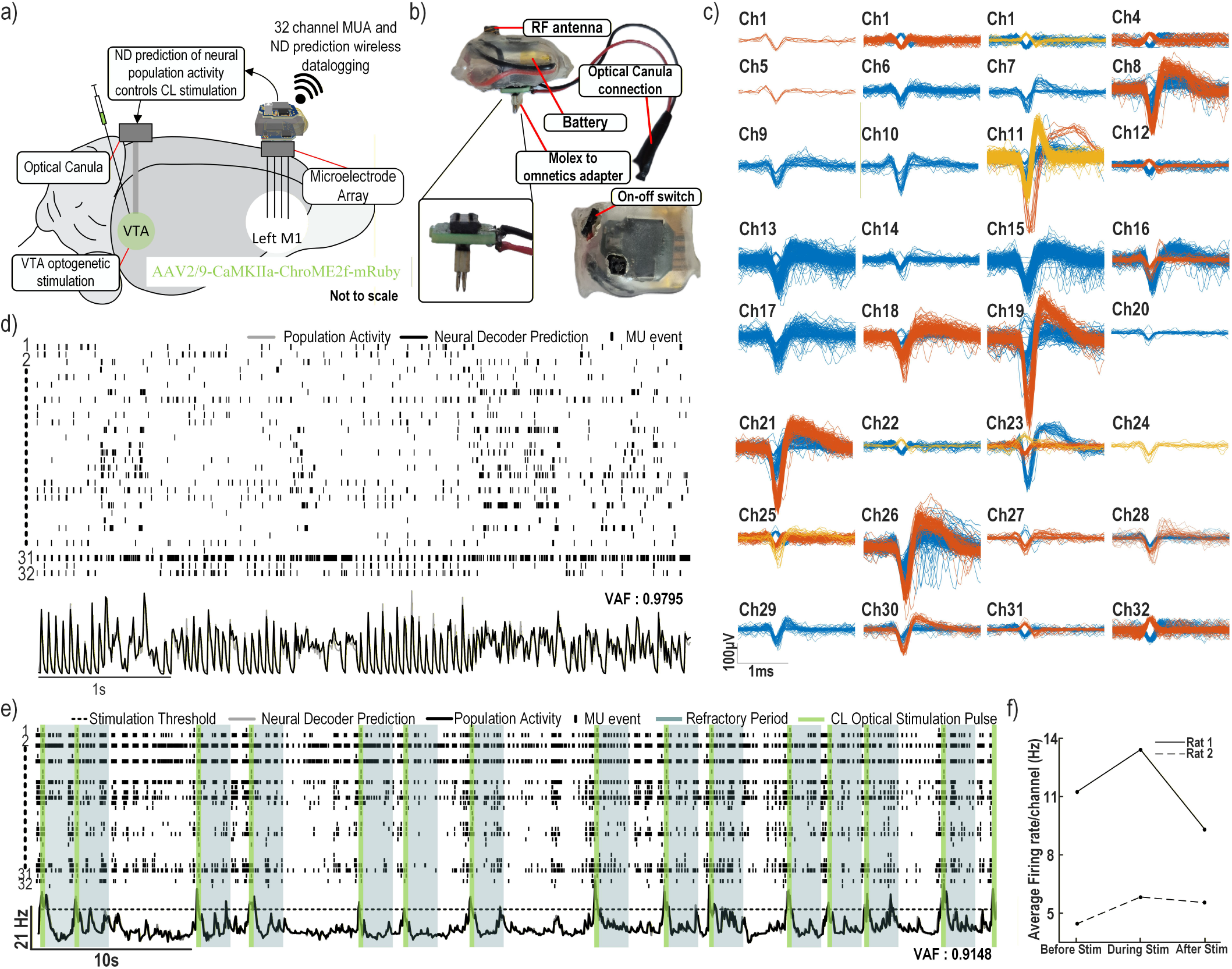
Neural decoding *in vivo* during anesthesia. a) Experimental setup with a 32-channel electrode array implanted in rat motor cortex and a fiber-optic cannula targeting the VTA. b) Packaged wireless headstage connected via a Molex-to-Omnetics adapter and coupled to the optical cannula for stimulation. Overlaid MUA from all 32 channels during anesthesia, showing representative waveforms. d) Raster plot with population firing rate and real-time decoder output. The decoder was trained on data from the same rat (10-ms bins, 10-ms LI decay) for fast, low-latency firing-rate prediction. e) Wireless closed-loop results using a longer accumulation model (50-ms bins, 150-ms LI decay), integrating larger temporal dynamics of the neural signal. When predicted activity exceeded the threshold (dotted line) for 50 ms, a 300-ms optical pulse was delivered, followed by a 2-s refractory period. Stimulation periods are shown in green, refractory period in grey. f) Average population firing rate across two rats before, during, and after closed-loop stimulation.

Figure 10(b) shows the packaged wireless system, with the battery and on–off switch positioned between the two PCB panels and protected with heat-shrink tubing. The headstage was connected to one of the two 32-electrode banks of the polymer array for the first rat, and to the electrodes of the MINT array for the second rat through the Omnetics connector on each probe. A custom adapter was used to interface the headstage *Molex* connector (0559090574) with the *Omnetics* A79026-001 (36 pins). In addition, a wired connection was made between the headstage and the optical cannula to control optogenetic stimulation through the wireless system (Figure 10(b)).

### C. Data Collection and Model Training

To train the wireless neural decoder, a multichannel AP recording was first acquired across all 32 electrodes to serve as the training dataset. The analog front-end bandpass filter was configured from 300 Hz to 5 kHz to capture the AP bandwidth of interest. To record 32 channels sampled at 20 kHz, the system uses its adaptive threshold detector together with the DWT compression core to transmit only the compressed APs and their timestamps to the base station, rather than the full raw signal, which would require a data rate of 10.24 Mbps and exceed the wireless link capacity (maximum 2 Mbps).

Multiple models can then be trained from the recorded AP waveforms and timestamps and wirelessly uploaded to the device to address different experimental objectives. As a proof of concept, two models were trained using the same 120 s recording. Both models were designed to predict a filtered version of the summed spike-count across all channels, serving as an estimate of the population firing rate.

The first model demonstrates the system’s ability to operate at very low latency using small accumulation windows. It uses 10 ms spike-count bins and a 10 ms leaky-integrator decay constant on all 32 channels, enabling rapid CL stimulation based on fast temporal dynamics.

The second model uses longer 50 ms bins and a 150 ms leaky-integrator decay constant to capture firing-rate dynamics over a longer timescale. For both models, PCA coefficients and an SVM regressor with a first-order polynomial kernel were trained to generate real-time population firing-rate predictions that can be used to trigger CL stimulation.

### D. Neural Decoder Results

The recorded MUA from the 32 electrodes under anesthesia was overlaid and clustered offline using PCA followed by k-means, enabling visualization of distinct neurons detected on each channel. The resulting waveforms are shown in Figure 10(c). Clear AP shapes are visible on multiple channels, with amplitudes reaching up to 400 *µ*V (e.g., channel 19), and several channels exhibit multiple distinct waveform clusters. These observations highlight the recording quality of both the system and the electrode arrays.

For the short-accumulation model reflecting faster temporal dynamics (Section VII-C), the live prediction generated by the headstage is shown alongside the actual population activity derived from the wirelessly logged MUA in Figure 10(d). A synchronized raster plot is also provided to allow qualitative comparison between the detected spikes and the predicted population firing rate. The raster is constructed from the logged timestamps, where each vertical line indicates a detected AP and each row corresponds to a different channel (with their associated AP waveforms shown in Figure 10(c)).

Population activity was computed by binning the logged timestamps into 10 ms windows, summing events across all 32 channels, and applying an averaging filter to smooth the resulting trace. The live on-device prediction closely matches this computed population activity, yielding a VAF of 0.9795. This strong agreement demonstrates that the model is well fitted and confirms the system’s ability to deliver accurate, low-latency predictions in real time.

### E. Closed-Loop Optogenetic Stimulation

For the CL experiment, a longer accumulation model (50 ms bin size, 150 ms leaky-integrator constant) was used together with a threshold-based rule: stimulation was triggered whenever the predicted population activity exceeded a predefined threshold for at least one bin (50 ms). A 2 s refractory period was imposed to prevent overstimulation. All parameters, including the threshold, timing, stimulation pulse parameters, and the neural decoder model itself, are user-defined and can be reprogrammed wirelessly during the experiment.

Figure 10(e) shows the spike raster plot alongside the model’s predicted population activity. The threshold is indicated by a dotted line, green rectangles mark the closed-loop triggered optical stimulation pulses delivered through the fiber-optic cannula, and gray rectangles indicate the refractory periods following stimulation. Each stimulation event consisted of a 50 mA, 300 ms light pulse. The model generated stimulation events that aligned with periods of sustained high firing across the recorded neural population.

The system reliably triggered stimulation during high-activity episodes, as shown in Figure 10(e). The VAF between the predicted population activity generated by the complete wireless neural decoder chain (neuromorphic feature extraction and SVM prediction) and the population activity computed offline from the wirelessly logged APs was 0.9148. This result further validates the wireless neural decoder, demonstrating its reprogrammability, precision, and suitability for autonomous CL stimulation.

Figure 10(f) shows the average population activity before, during, and after the CL stimulation experiment for each rat, revealing an increase in average population activity in the motor cortex during the VTA stimulation experiment in both cases, which suggests that VTA stimulation modulates motor cortex activity and is associated with an increase in population firing rate under these experimental conditions.

Overall, these results demonstrate that the platform can perform real-time wireless neural decoding and closed-loop stimulation based on neuromorphic feature extraction and SVM-driven population activity estimation, with fully reprogrammable parameters across all 32 neural channels.

## VIII. Discussion

The proposed wireless headstage demonstrates high-performance, low-latency neural decoding validated on both datasets and *in vivo*. On the rhesus macaque center-out task, the decoder achieved *R*^2^ = 0.85 for Y-axis and 0.66 for X-axis forces using six PCA components derived from 32-channel MUA with leaky integration (Table I). SVM model clustering reduced support vectors, decreasing model size from 80 kb to 22 kb without loss of accuracy, or improving *R*^2^ by 0.09–0.17 when constrained to 15–20 kb (Figure 4). In rats performing a lever-pull task, the decoder generalized to binary classification with high accuracy across 16 channels (Table 9). *In vivo*, the headstage accurately predicted population activity (VAF = 0.9795 for the 10-ms bin model; 0.9148 for the closed-loop stimulation model) and reliably triggered optical pulses aligned with high-activity episodes (Figure 10(e-f)).

Compared with existing neural decoders, the proposed headstage achieves a combination of flexibility, computational efficiency, low latency enabling timing-sensitive closed-loop control, and *in vivo* validated performance not simultaneously demonstrated in prior work. Table I shows that the feature–model pipeline (binned MUA, leaky integrator, PCA, NL-SVM) maintains high decoding accuracy while reducing feature dimensionality by more than an order of magnitude. Using six PCA components, the decoder achieves *R*^2^ comparable to CNN- or SVM-based models operating on feature vectors 50 times larger. Temporal compression via the leaky integrator and spatial compression via PCA enable real-time inference on a resource-limited FPGA without the memory footprint required by conventional deep or recurrent models. This allows closed-loop interventions within the 10–20 ms timescale relevant for timing-sensitive plasticity mechanisms, such as STDP.

At the system level (Table III), prior FPGA- or ASIC-based implementations typically process only spikes [51], perform neural decoding without closed-loop capability or without a complete wireless platform [41], or rely on tethered PC-based decoding [15], [19], [28]. Many high-performance decoders require *>*100 kB memory, hundreds of MAC units, or multi-millisecond latency, limiting deployment on lightweight headstages and precluding real-time interventions that could influence neural dynamics in a biologically meaningful manner for some applications. In contrast, our implementation requires only three MAC units and ≤ 10 kB model memory to deliver sub-millisecond latency, physiologically relevant closed-loop modulation based on 32-channel MUA extraction and on-device PCA, with wireless on-the-fly reprogramming capability. Unlike many prior FPGA or IC-based decoders, our platform demonstrates full *in vivo* closed-loop operation with real-time prediction, wireless data logging, and optical stimulation triggered by decoded neural activity.

**TABLE III.** System-level comparison with state-of-the-art neural decoders and signal processing systems.

|  | *Bin Altaf*<br>2016, [50] | *Wang*<br>2019, [51] | *Abbasi*<br>2018, [19] | *Barroso*<br>2019, [15] | *Bouton*<br>2016, [16] | *Katoozian*<br>2020, [42] | *Katoozian*<br>2024, [41] | *Gagnon-Turcotte*<br>2019, [5] | *Oh 2025,*<br>[28] | **This paper** |
| --- | --- | --- | --- | --- | --- | --- | --- | --- | --- | --- |
| **Application** | Seizure Detector | Spike Sorter | Neural Decoder | Neural Decoder | Neural Decoder | Neural Decoder | Neural Decoder | Spike Sorter | Neural decoder | Neural decoder / Classifier |
| **Signal of interest** | EEG | Spike | Spike | Spike | Spike, LFP, MUA | Spike | Spike | Spike | LFP / accelerometer | MUA |
| **Features** | FFT | N/A | Firing rate | Binned spike counts | WT | Binned counts | Binned counts | Firing rate | Time–frequency content | Binned counts |
| **Time window (ms)** | 2000 | N/A | 100 | 50 (30 windows history) | 100 | 35 (30 windows history) | 10-70 | N/A | 4000 | **5-50** |
| **Decoder type** | NL-SVM | Template Matching | Threshold | LSTM | NL-SVM | SVM | HDC | Vector based | CNN | NL-SVM |
| **Decoding Platform** | On-device (IC) | Host (FPGA) | Host (PC) | Host (PC) | Host (PC) | On-device (FPGA) | On-device (FPGA) | On-device (IC) | Host (PC) | **On-device (FPGA)** |
| **Wireless** | No | No | No | No | No | No | No | Yes | Yes | **Yes** |
| **Latency (ms)** | 2000 | 2 | 12.3 | N/A | 800 | 180 | - | 0.6 | N/A | 0.254 |
| **On the fly reprogramming** | No | No | No | N/A | No | N/A | N/A | Yes | Yes | **Yes** |
| **Stimulation** | Electrical | N/A | Optical | N/A | Electrical | N/A | N/A | Optical | N/A | **Optical** |
| **Nb. of Channel** | 8 | 1 | 32 | 32 | 96 | N/A | N/A | 10 | 32 | 32 |
| **Level of integration\*** | Partial (IC only) | Partial (No stim) | Yes | No | Yes | No | No | Yes | Partial (No Stim.) | **Yes** |
| **In vivo validation** | No | No | Yes | No | Yes | No | No | Yes | Yes | **Yes** |

\*Indicates whether the neural decoder is integrated into a bidirectional interface, enabling closed-loop neuromodulation based on decoder outputs within a practical autonomous system for live experiments.

Key contributions of the platform, uniquely integrated within a compact, battery-powered headstage, include:

- High-rate MUA-based feature extraction from 32 neural channels
- Neuromorphic temporal integration and dimensionality reduction via leaky integrator and on-device PCA
- Nonlinear SVM inference on a resource-constrained FPGA, achieving sub-millisecond latency for physiologically relevant closed-loop control
- Wireless reprogrammability of decoder parameters and stimulation settings
- *In vivo* closed-loop optogenetic stimulation triggered by decoded neural activity

Together, these capabilities enable a flexible, resource-efficient platform for real-time neural decoding and closed-loop neuromodulation in freely moving animals. This represents the first fully wireless headstage integrating neuromorphic feature extraction, FPGA-based decoding, and *in vivo* closed-loop stimulation, with latency compatible with timing-sensitive neural plasticity mechanisms.

## IX. Conclusion

We presented a compact, wireless neural headstage capable of real-time, closed-loop neural decoding and optogenetic stimulation in freely behaving rodents. The system was validated across multiple datasets and in *in vivo* experiments, where the system simultaneously recorded multi-channel M1 activity and autonomously delivered closed-loop optogenetic stimulation to the VTA in real time. These experiments demonstrated accurate, low-latency on-device predictions of population firing rate and reliable, temporally precise triggering of stimulation. Together, the results highlight the system’s ability to support real-time, feedback-driven neuromodulation through indirect dopaminergic reinforcement mechanisms.

By integrating multi-unit recordings, efficient feature extraction, dimensionality reduction, and optimized device decoding, the platform achieves high performance while operating within the memory and computational constraints of a small FPGA. Its wireless reprogrammability allows flexible model deployment and stimulation parameter adjustment during experiments, supporting a wide range of behavioral paradigms. This work demonstrates a fully self-contained, versatile neural interface capable of supporting advanced closed-loop neuroscience experiments in naturalistic settings, opening new opportunities for probing circuit dynamics, studying disease mechanisms, and developing adaptive neuromodulation therapies in small-animal models.

## Acknowledgment

We acknowledge support from the Canada Research Chair in Smart Biomedical Microsystems.

